# Exploring the interaction between immune cells in the testicular microenvironment of azoospermia combining RNA-seq and scRNA-seq

**DOI:** 10.1101/2022.12.12.520033

**Authors:** YuanYuan Wu, JinGe Huang, Nan Ding, MengHao Lu, Fang Wang

**Author notes:** Correspondence: Fang Wang, E-mail addresses.

## Abstract

Non-obstructive azoospermia is the most serious cause of male infertility. The testis has a special immunological environment, but the relationship between immune cells in the testicular microenvironment is still unclear. Therefore, it is urgent to identify the interaction mechanism and molecular determinants of immune cells in the testicular microenvironment. To further elucidate the etiology of azoospermia and provide a reference for the treatment of azoospermia. The GSE145467 and GSE9210 datasets were analyzed by Limma package, and then the differential genes were analyzed by enrichment analysis and protein-protein interaction analysis. In addition, we combined single-cell analysis(scRNA) to identify immune cell types and verified the expression of Hub genes in these immune cells. Finally, CellChat was used for cell-to-cell communication analysis. We found the distribution of immune cells in the microenvironment of Y chromosome AZF region microdeletions (AZFa_Del), idiopathic NOA (iNOA), and Klinefelter syndrome (KS) was significantly different from that of normal adults, especially monocytes/macrophages. In normal subjects, monocytes/macrophages mainly played the role of the signal source, while in patients with azoospermia, monocytes/macrophages mainly received signals from other immune cells. Monocytes/macrophages in AZFa_Del, iNOA, and KS communicated with other immune cells mainly through MDK-LRP1, PTN-NCL, and MDK-NCL ligand-receptor pairs respectively. Our research provides new ideas for the pathogenesis and treatment of azoospermia.

**Author Summary:** This article uses the datasets in the public database, including RNA-seq and scRNA-seq. It is a secondary analysis of these databases. Previous studies have found the destruction of Sertoli cells in the testicular microenvironment using scRNA datasets. We have analyzed immune cells in the testicular microenvironment based on previous studies. Found huge changes in macrophages and abnormal macrophages communication through cell communication analysis, The receptor-ligand pairs were screened to provide a basis for the study of macrophages in the testicular microenvironment and the treatment of azoospermia.

## Introduction

Infertility can destroy a family. 15% of couples in the world are suffering from infertility. Male factors account for about 50%, the main reason is abnormal spermatogenesis(1). Nonobstructive azoospermia (NOA) is the most serious cause of male infertility(2), the main etiology is Klinefelter syndrome (KS), Y chromosome AZF region microdeletions (AZF_Del), and idiopathic NOA (iNOA) (3). The latest research has found severe damage to the testicular microenvironment Sertoli cells in NOA patients, Abnormal immune responses have been found in patients with KS and AZFa_Del (4). Testicular immune privilege protects immunogenic germ cells from systemic immune attack(5). Testicular macrophage (TM) act multiple roles in maintaining normal testicular function and homeostasis(6). The research has found that when macrophages were damaged, the testosterone level, sperm counts, and sexual desire of mice would be significantly reduced (7). TM also has played a relevant role in the morphogenesis of the testicular vasculature in testis cord formation and clearance of misguided germ cells in the mesonephros on their way to the gonad by phagocytosis(8). While in other research has found that enhanced numbers of macrophages correlated with fibrotic remodeling in the testicular interstitial space and impaired permeability of the blood-testis barrier in the Aix^-/-^/Mer^-/-^mouse(9). In this study, we aim to analyze the relationship between immune cells in the testicular microenvironment in azoospermia and explore the basis of their communication to provide ideas for the pathogenesis and future treatment of azoospermia.

With the rapid development of high-throughput technologies, bulk RNA sequencing (RNA-seq) technologies have been used widely in gene expression research at the population level. In recent years, single-cell sequencing (scRNA-seq) technologies have also provided the possibility to explore gene expression profiles at the single-cell level. Single-cell sequencing allows high-throughput sequencing analysis of the genome, transcriptome, and epigenome of individual cells, reflecting intercellular heterogeneity, and gene and molecular functional diversity(10). Jin et al. developed CellChat, a tool capable of quantitatively inferring and analyzing intercellular communication networks from single-cell RNA sequencing data (11). Cellular communication is the process by which cells receive, handle, and transmit signals from other surrounding cells or themselves which plays an important role in coordinating various biological processes.

In this study, we constructed a protein-protein interaction PPI network for the differential genes in GEO datasets GSE145467 and GSE9210. GO and KEGG analysis was used on the differential genes. We found that azoospermia was associated with immune disorders. Cluster analysis on single cell dataset GSE149512 showed that macrophages were mainly in NOA, and CellChat analysis showed that macrophages in the immune microenvironment of AZFa_Del, iNOA, and KS were mainly as acceptors. The ligand-receptor pairs that affect the communication between macrophages and other cells are MDK-NCL in iNOA, MDK-LRP1 in AZFa_Del, and MDK-NCL in KS. These ligand-receptor pairs may be future targets for the immunotherapy of azoospermia.

## Results

### Identification of DGEs

GEO2R analysis identified 4099 DEGs in GSE145467 and 673 DEGs in GSE9210. Used ggplot2 software package to draw volcano map in R (**Fig 1A**). We obtained 168 upregulated genes and 49 downregulated genes, as shown in the Venn plot (**Fig 1B**).

**Figure 1.**
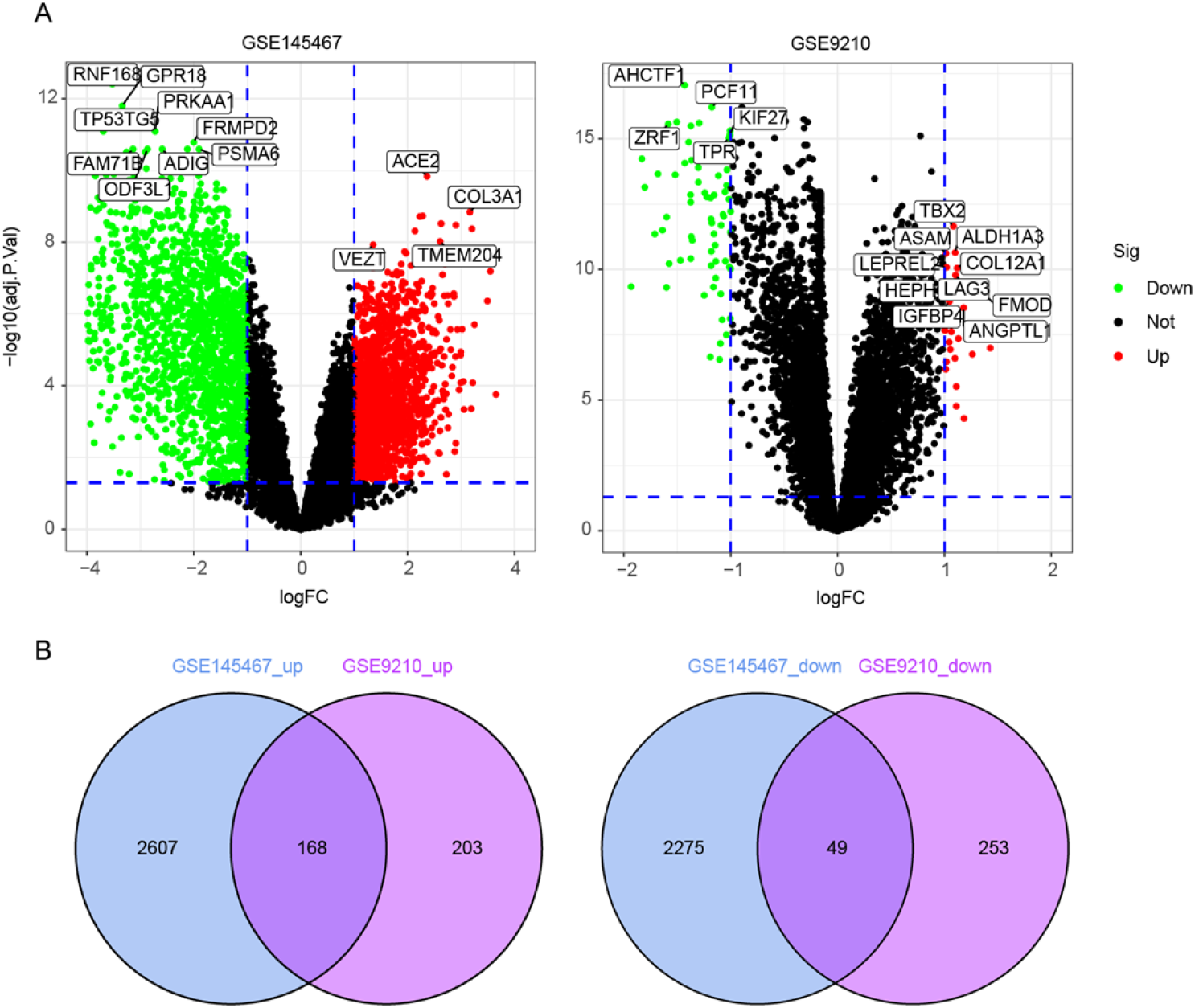
Screening results of DEGs. (A) A volcano map of DEGs in the GSE 9210 and GSE 145467 datasets. The red, green and black points represent genes that were upregulated, downregulated and showed no significant differences in expression, respectively. (B) A Venn diagram displaying the number of DEGs in the two datasets from the GEO database.

### GO enrichment analysis and KEGG pathway analysis of the DEGs showed functional enrichment in immune regulation

To explore the potential signaling pathways and biological functions involving the common DEGs, we used the DAVID database to perform GO annotation and KEGG pathway enrichment analyses, which were visualized using ggplot2 to produce bubble plots (P<0.05) (**Fig 2**). GO analysis identified that the DEGs are mainly associated with biological processes such as immunoglobulin mediated immune response, lymphocyte mediated immunity, and B cell mediated immunity. In molecular function, the main changes focused on amide binding, peptide binding, and glycosaminoglycan binding. The changes in cellular components were mostly collagen-containing extracellular matrix, blood microparticle, and MHC protein complex (**Fig 2A**). (**Fig 2B**) shows that the top five KEGG-enriched pathways associated with the DEGs were staphylococcus aureus infection, phagosome, systemic lupus erythematosus, Tuberculosis, steroid hormone biosynthesis, and antigen processing and presentation signaling pathways. These results showed that DEGs play an important role in NOA patients, which is related to the activation status of immune cells in TM of NOA patients.

**Figure 2.**
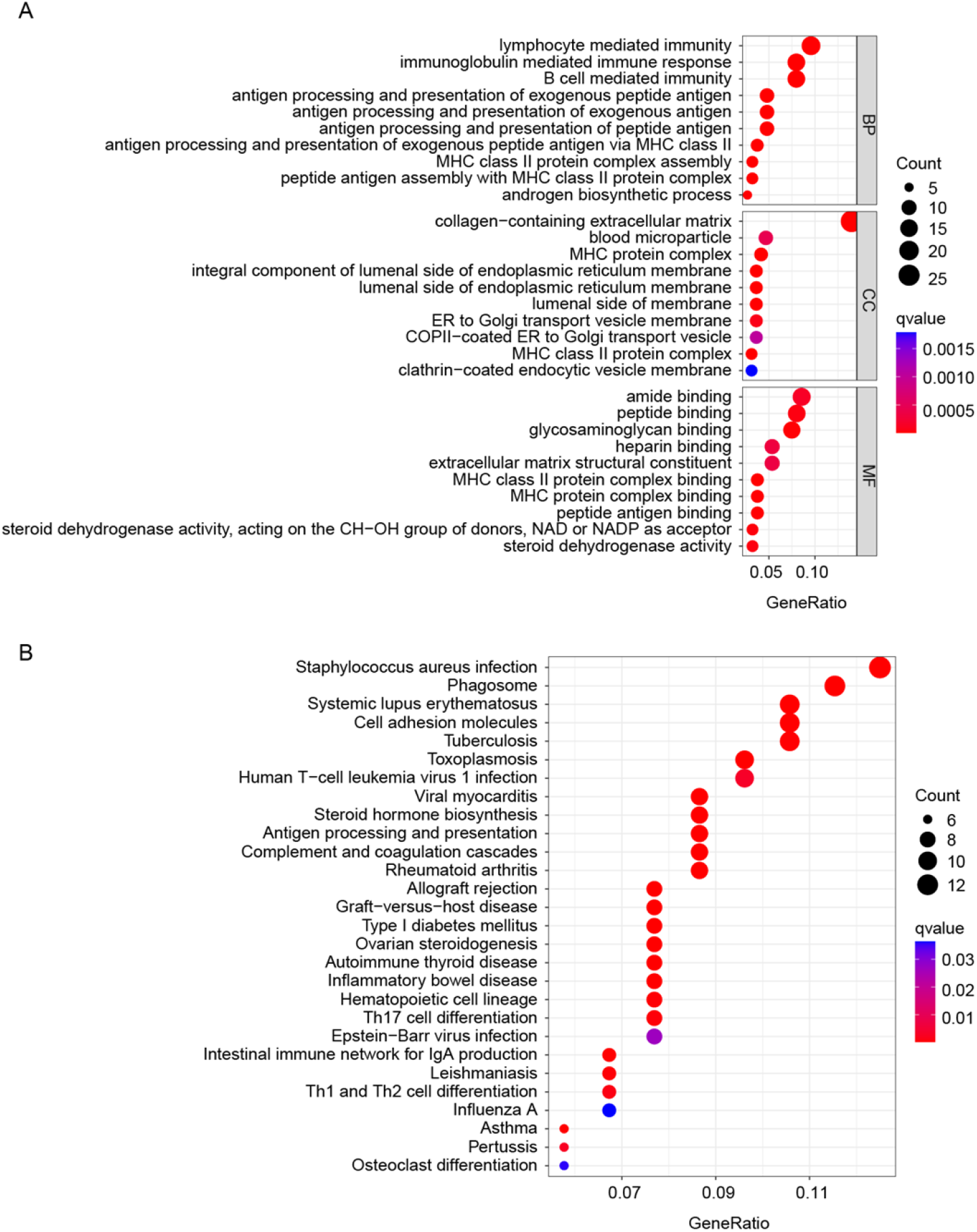
GO and KEGG pathway enrichment analysis of the DEGs. (A) GO analysis of DEGs, including biological process, cell components, and molecular functions. (B) KEGG pathway enrichment analysis of the DEGs. The GO analysis and KEGG pathway enrichment analysis of the DEGs were completed using DAVID and visualized using the ggplot2 package.

### PPI network construction and the identification of hub gene

To further investigate the interaction of the identified DEGs, we constructed a Protein-protein interactions (PPI) network using STRING and Cytoscape to explore the interactions and central genes of DEGs. A total of 217 DEGs were uploaded to the STRING website and the comprehensive score >0.4 was used as the cut-off standard. First, the PPI network was constructed using Cytoscape (**Fig 3A**) which included 139 nodes and 360 edges. Nodes represent proteins, edges represent interactions between proteins, and the number of edges connected by genes is positively correlated with the importance of their functions in the PPI network. The top 15 hub genes with high connectivity in the PPI network were identified using the plug-in maximal clique centrality (MCC) of cytohubba: CXCL12, CFH, FMOD, AR, DCN, HSP90AA1, C1QA, CSF1R, TIMP3, CTSK, IGF1, APOE, C1QB, COL3A1, TYROBP. All those genes were upregulated, except HSP90AA1 (**Fig 3B**).

**Figure 3.**
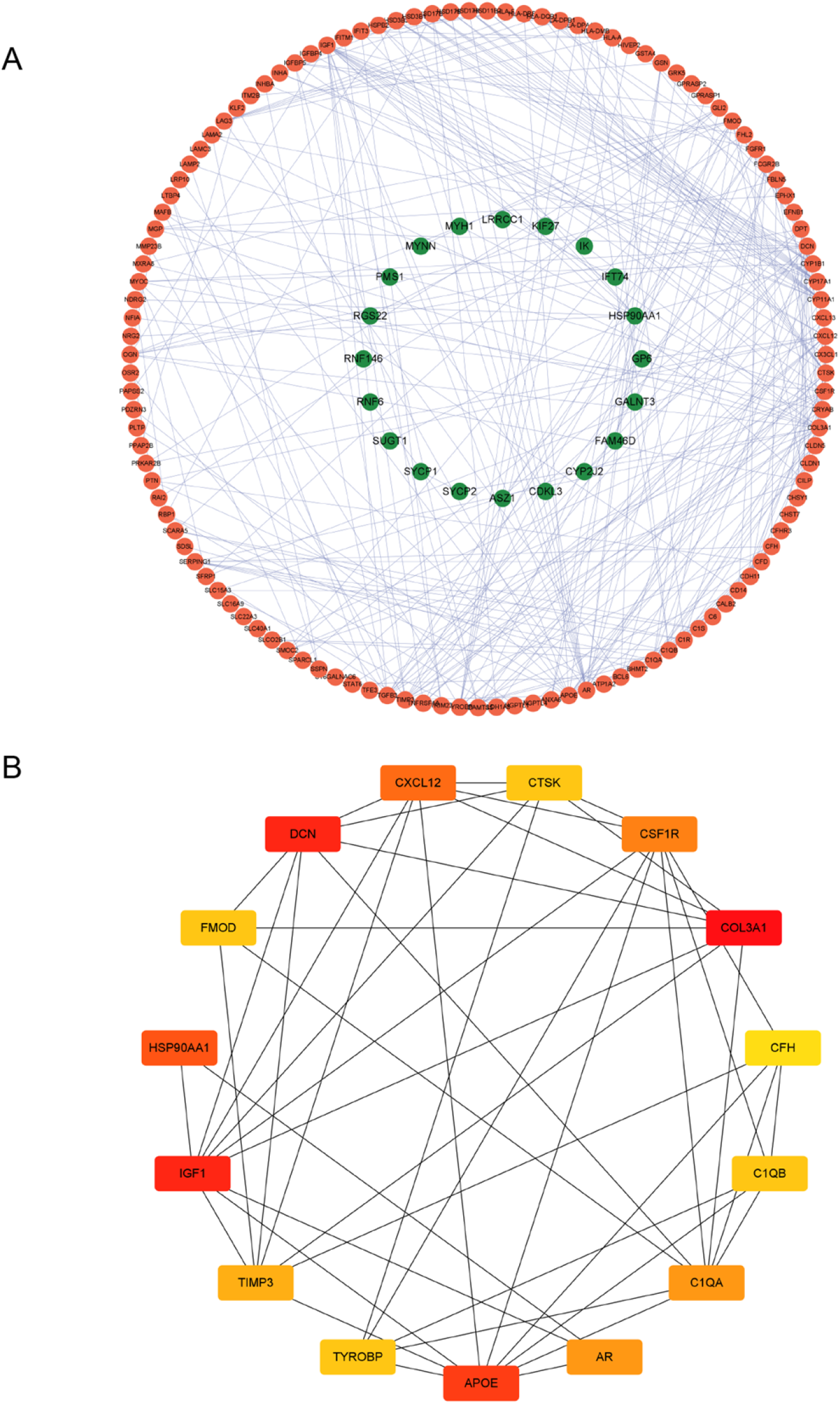
Construction of a PPI network and the identification of hub gens. (A) PPI network of DEGs, the outer circle represents upregulated genes, and the inner circle represents downregulated genes. (B) Top 15 DEGs obtained by MCC of cytohubba.

### Immune cells in NOA and normal samples

We used the CIBERSORT package to analyze immune cell infiltration in the dataset, results showed that compared with the normal group, CD4^+^T, resting NK, activated dendritic, and activated mast cells were lower in the NOA group, while monocytes, CD8^+^T, resting dendritic, resting mast cells were significantly higher (P<0.05), as shown in (**Fig 4**).

**Figure 4.**
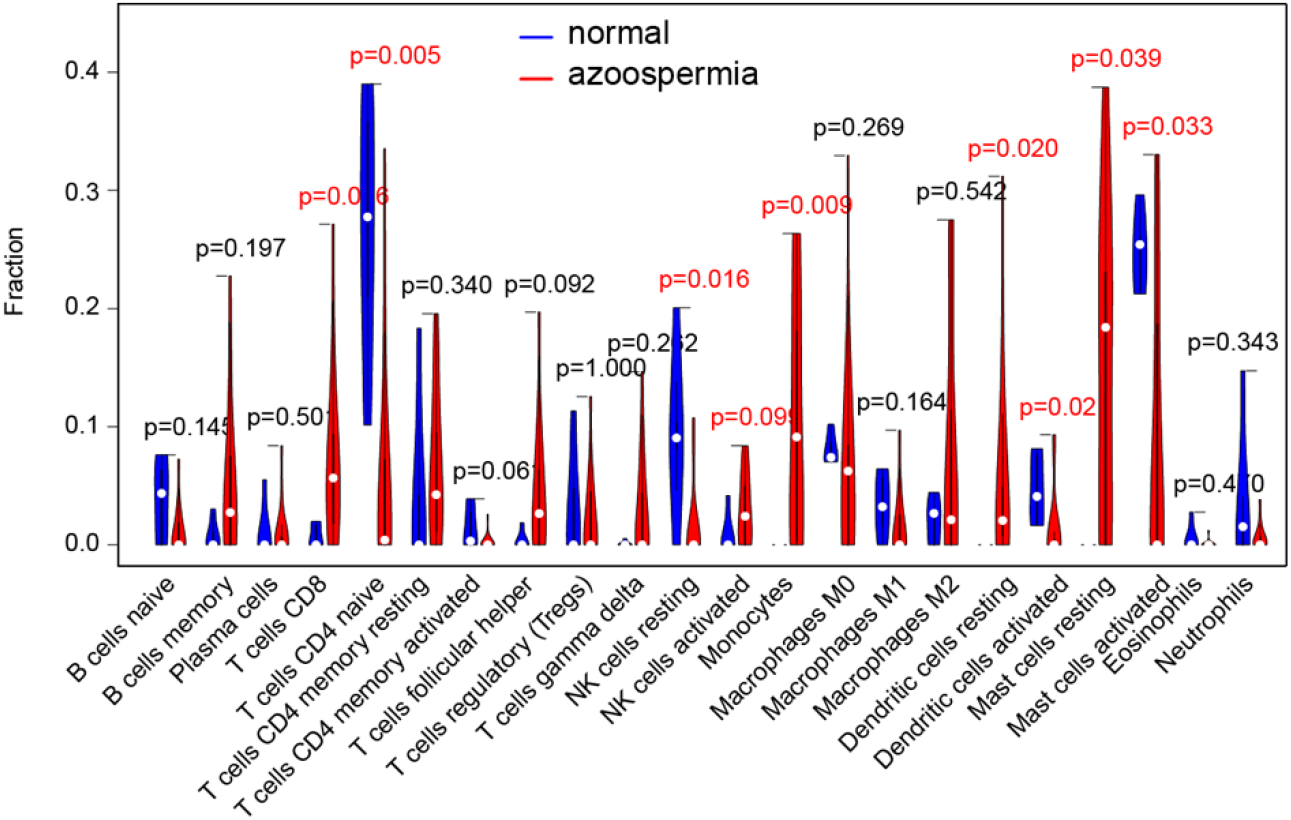
Immune cells in NOA and normal samples. Difference of immune cell expression between azoospermia and normal

### ScRNA-seq analysis and hub genes in immune cells

Firstly, 10 healthy subjects in the scRNA-seq data of GSE149512 were divided into 4 stages according to their growth and development stages (**Fig 5B**), analyzed, and identified. Used a dimensionality reduction algorithm (t-SNE) to visualize the results. The types of immune cells in different periods of normal testicular microenvironment were identified, mainly including T cells, B cells, and monocytes/macrophages (**Fig 5A**). We found that monocytes/macrophages differ in different age groups. To further understand the differences between the testicular cells of normal subjects and patients with NOA, cluster analysis was used in AZFa_Del, KS, and iNOA (**Fig 5A**), monocytes/macrophages between normal adult and three types of NOA showed the greatest dissimilarity than others immune cell. Combined with the hub genes, we further analyzed the expression of hub genes in the immune cells of NOA and normal adults (**Fig 6**). The hub genes were highly expressed in most monocytes/macrophages, In AZFa_Del patients, about 75% of monocytes/macrophages highly expressed DCN and APOE (**Fig 6**). In iNOA patients, the expression of hub genes in monocytes/macrophages was surprisingly similar to normal adults (**Fig 6**). In KS patients, about 75% of monocytes/macrophages and CD4^+^T cells expressed DCN, APOE, TIMP3, and about 75% of CD4^+^T cells highly expressed IGF1 (**Fig 6**). Our results suggest that the changes in the testicular immune microenvironment in NOA patients are mainly in monocytes/macrophages. Therefore, studying the relationship between macrophages and other immune cells in TM is helpful to discuss the pathogenesis of azoospermia and provide evidence for treatment.

**Figure 5.**
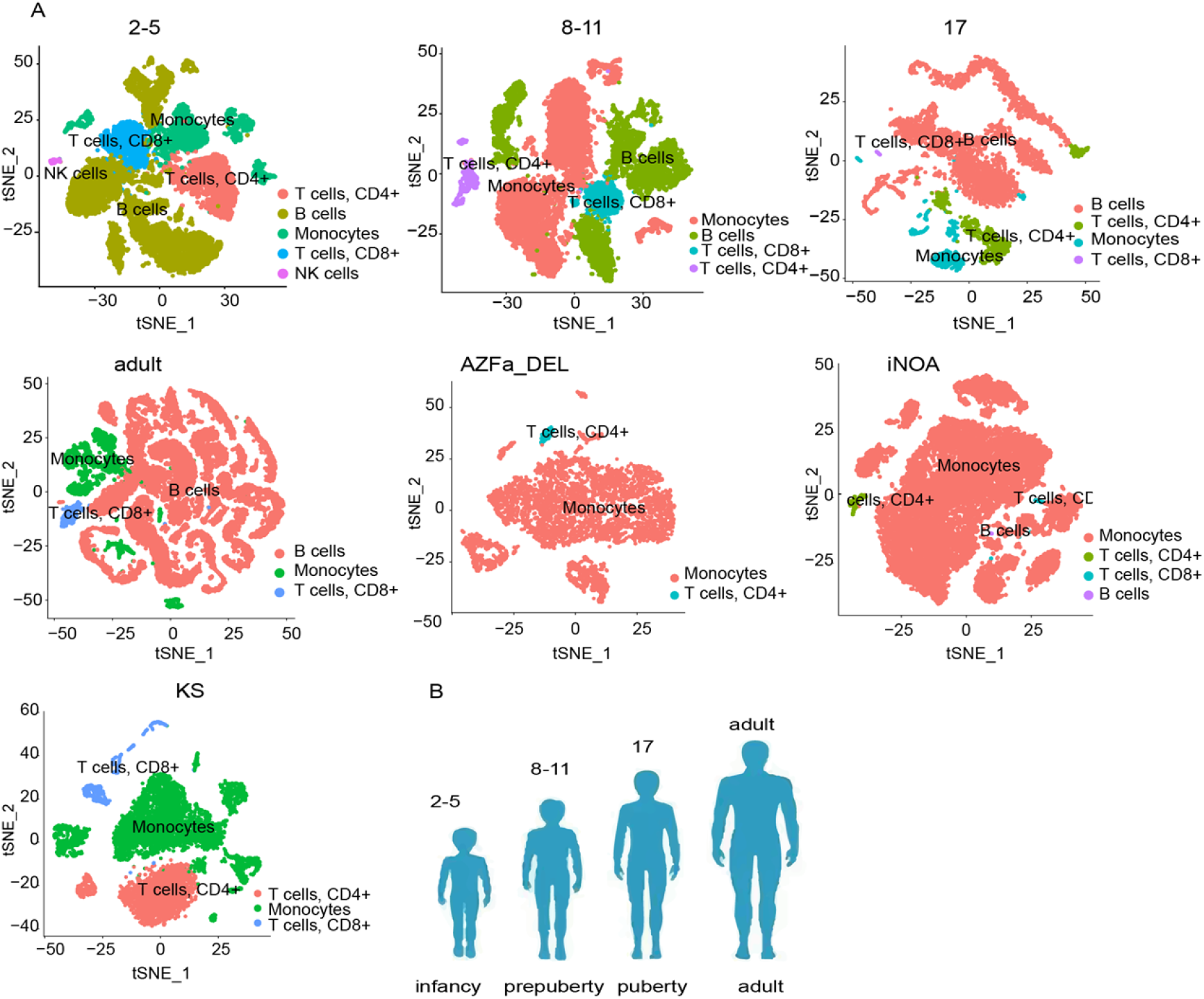
Immune cell types in testicular microenvironment were identified by scRNA-seq. (A) Cluster analysis of testicular immune cells in normal and NOA subjects, and the results were visualized by t-SNE dimensionality reduction clustering. (B) 10 normal control groups were divided into 4 groups according to growth and development stages.

**Figure 6.**
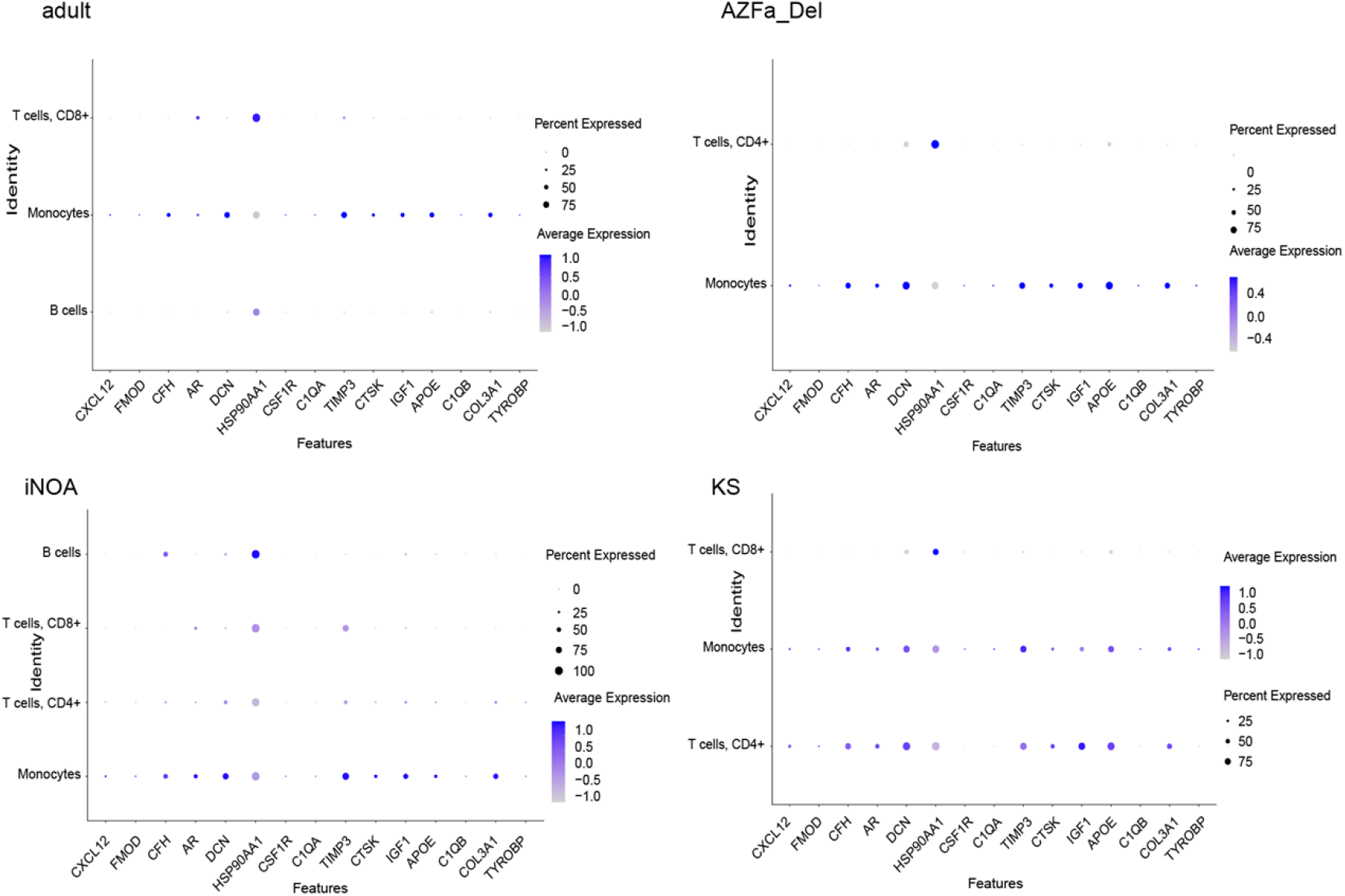
The expression of hub genes in different immune cells. The differences of hub genes expression between normal adults and three kinds of azoospermia were compared

### Integrated analysis reveals the basis of the interaction between immune cells in the testicular microenvironment

To identify the potential interaction**s** between different immune cells, we used CellChat analysis on a dataset from the GEO database (GSE149512). CellChat contains a database of receptor-ligand interactions containing 2,021 verified molecular interactions. CellChat can identify the key features of cell-to-cell communication in a given scRNA-seq data set and predict potential signaling pathways that are currently less studied. To obtain more critical cell-cell interactions in the testicular microenvironment, we selected the receptor-ligand related pathway with the largest possibility among immune cells and the smallest P value, and then calculated and visualized the contribution of each receptor ligand to the whole signal pathway.

### Normal subjects

To investigate the complex signaling networks of immune cells in the testicular microenvironment, we performed an unbiased ligand-receptor interaction analysis between these testicular cell subsets by CellChatDB(12). In infancy, prepuberty, puberty, and adult, we found 22, 19, 25, and 16 interactions between macrophage ligands and receptors from other immune cells, and 9, 12, 21, and 7 interactions between ligands from other cells and macrophage receptors, respectively (**Figs 8A** and **S1 Fig**). Obviously, the former was more than the latter. Different stages have different cell types and interaction characteristics. Focusing on the interaction between different immune cells, we found that the ligand-receptor pair related to the MIF pathway in infancy has the greatest possibility and the minimum P value (**Fig 7**), and the MIF-(CD74+CXCR4), MIF-(CD74+CD44) ligand-receptor pair has a good contribution to the MIF pathway (**Fig 8B**). While in prepuberty, puberty, and adult, the ligand-receptor pair related to the PTN pathway has the largest possibility and the smallest P value (**Fig 7**), PTN-NCL has the strongest contribution to the PTN pathway (**Fig 8B**). In our study, PTN-NCL was involved in four groups. This indicates that the PTN pathway was of great significance to the connection of immune cells in human growth. Interestingly, NK cells in infancy acted as signal receptors to receive signals from other cells including itself through MIF-(CD74+CXCR4), MIF- (CD74+CD44), and macrophages only played the role of signal source (**Fig 8C**). In prepuberty, and puberty, macrophage played the role of the signal source to affect other immune cells through PTN-NCL (**Fig 8C**). In the adult, macrophages acted as signal sources and receptors to communicate with other cells through PTN-NCL (**Fig 8C**). NCL is expressed in every immune cell at different stages. whereas PTN is mainly expressed in macrophages (**Fig 8.D**).

**Figure 7.**
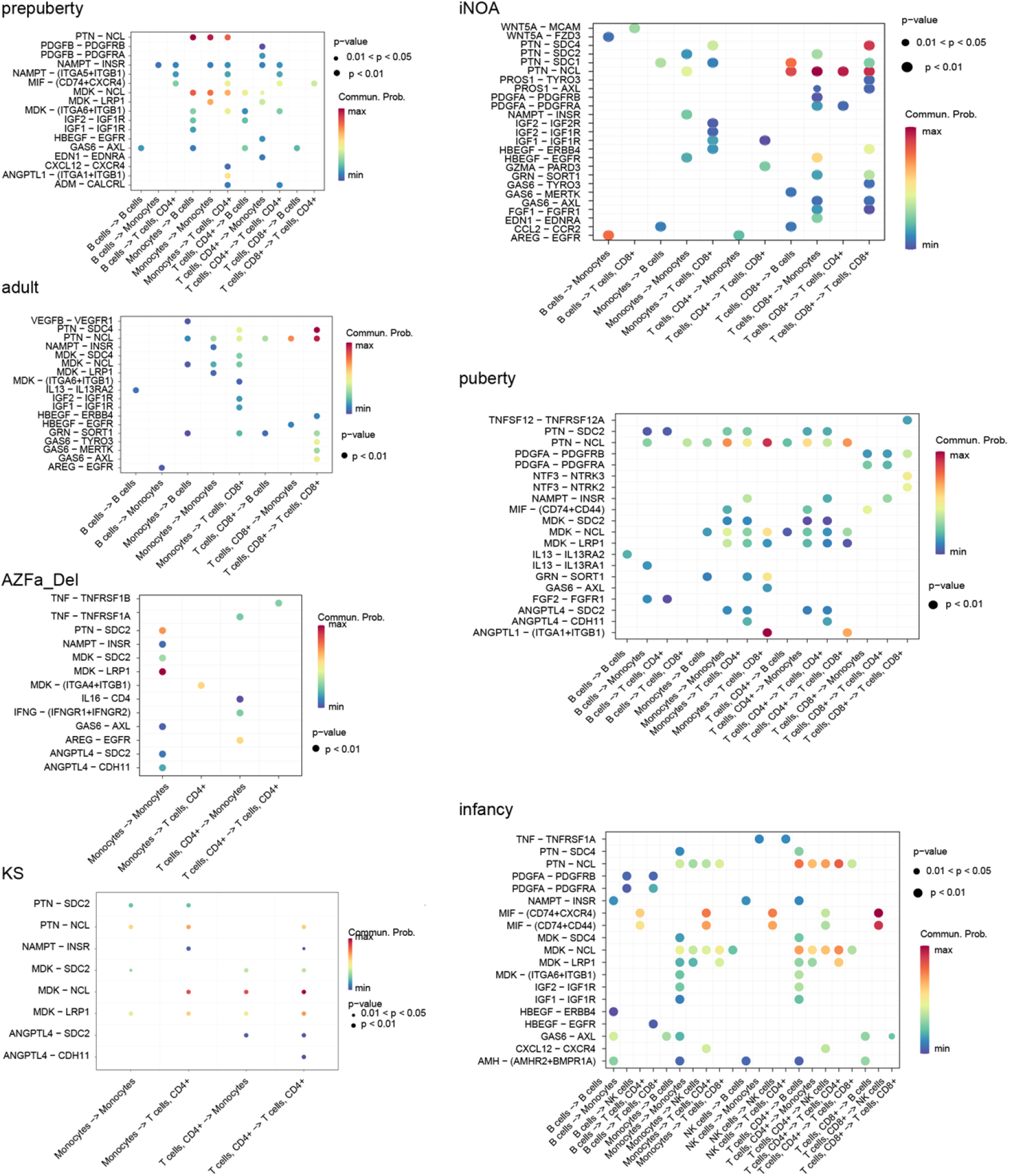
Bubble diagram of receptor ligand pair in interaction of immune cells, bubble size represents P value and color represents possibility.

**Figure 8.**
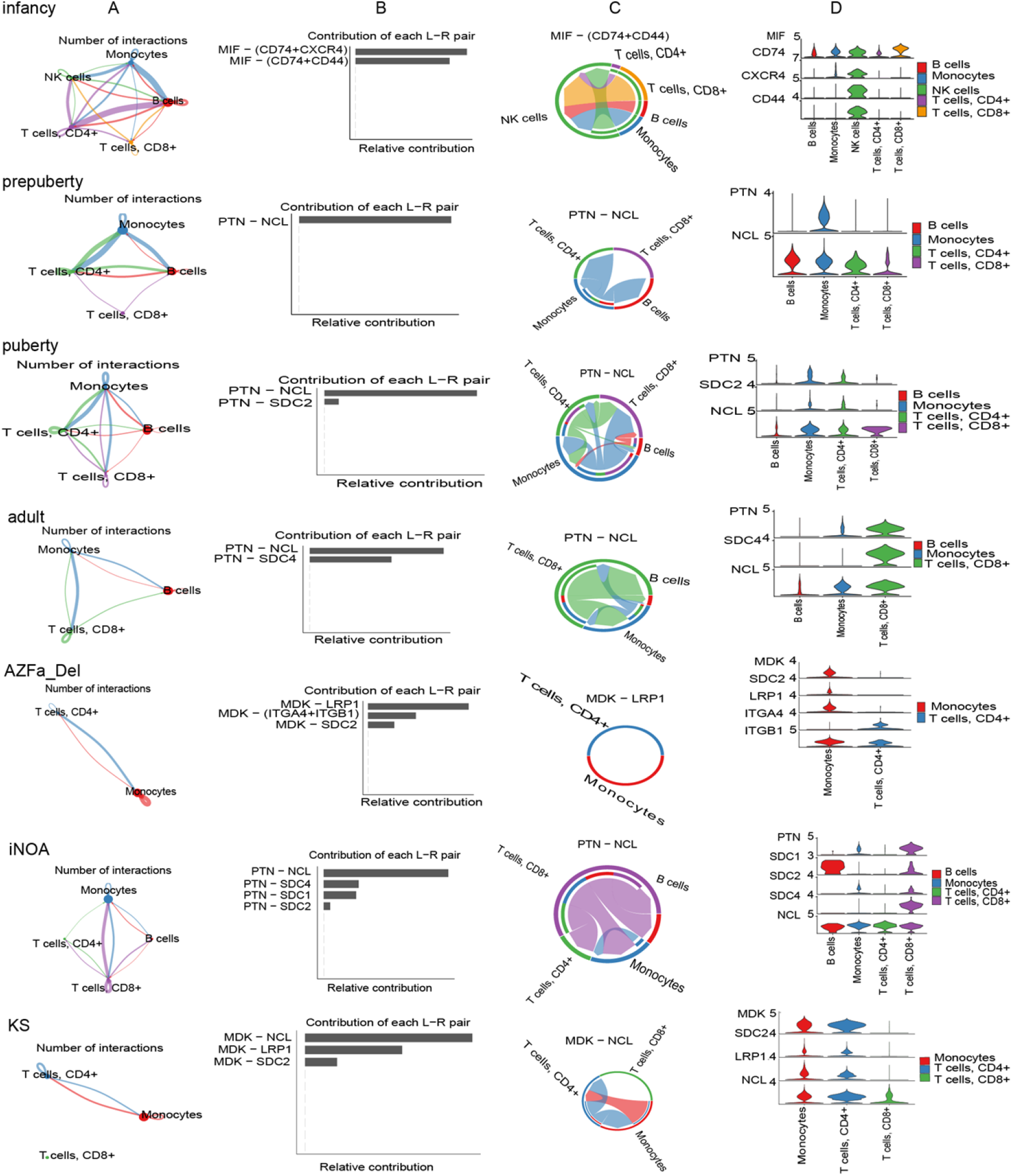
Cell communication network diagram, (A) the number of ligand receptor interactions detected between different cell types, (B)ligand receptor contribution to the overall signaling pathway, (C) receptor ligand pair interactions between immune cells and the distribution and (D) expression level of signal gens involved in the signal pathway networks. We compared the difference of immune cell signal communication between normal subjects and three kinds of azoospermia.

### NOA patient

Similarly, CellChatDB performed unbiased ligand-receptor interaction analysis on NOA testicular immune cells. In AZFa_Del, iNOA and KS, we found 8, 12, and 9 interactions between macrophage ligands and receptors from other immune cells, and 10, 17, and 8 interactions between ligands from other cells and macrophage receptors respectively (**Figs 8A**, **S1 Fig**). The latter was more than the former which is obviously opposite to the normal adult group. This suggested that abnormal spermatogenesis might be related to the over-activation of macrophages. Like normal adults, PTN-NCL related to the PTN pathway played an important role in iNOA patients (**Fig 7, 8B**). Unlike normal people, macrophages could only receive signals from themselves and CD8^+^T cells through PTN-NCL (**Fig 8C**), Although the NCL receptor was also expressed in CD8^+^T cells (**Fig 8D**). iNOA patients had similar types of immune cells to adults. But their distribution of various cells is greatly different from normal people. AZFa_Del patient macrophages could only receive signals from themselves through MDK-LRP1 (**Figs 7, and 8C**) and the LRP1 receptor was only expressed in macrophages (**Fig 8D**). KS patients not only could play the role of the signal source, but also received signal from CD4^+^T through MDK-NCL (**Figs 7, and 8C**). NCL is expressed in CD8^+^T cells, but CD8^+^T cells do not communicate with other cells (**Fig 8D**). Our results showed that macrophages communicate differently with other cells in NOA and normal adults. MDK-LRP1, PTN-NCL, and MDK-NCL ligand-receptor were key points between macrophages and other immune cells in AZFa_Del, iNOA, and KS patients respectively.

## Discussion

When patients learned that they suffered from azoospermia, they were very depressed. They would ask us what caused azoospermia. We could only explain the problems of chromosomes and genes. However, some patients with azoospermia could find sperm through the testicular puncture. The research found that a total of 211 patients had maturation arrests, and the overall sperm retrieval rate was 52%(13). This shows that some patients are capable of spermatogenesis. In addition to their poor spermatogenesis, the attack of immune cells on sperm may also be an important reason for azoospermia. Nowadays, there is still no effective method for azoospermia, which require us to study its mechanism more widely, explore new treatment methods, and cooperate with existing treatment methods to bring hope to patients. It has been found that Sertoli cells in the testicular microenvironment of patients with NOA were severely damaged(3), This is a remarkable discovery, but few studies have been conducted on the influence of immune cells in the testicular microenvironment on spermatogenesis. In this study, we combined Bulk RNA-seq and scRNA-seq. Bulk RNA-seq measures the average transcript level of a cell population, which can quickly identify biological markers of disease. scRNA-seq technology can generate expression profiles of individual cells for analysis of heterogeneous cell populations and identification of cell types. The understanding of the phenotype of immune cells in the testicular microenvironment is crucial to understanding the immune mechanism of azoospermia.

We identified the intersection of DEGs between azoospermia and normal samples by analyzing datasets GSE145467 and GSE9210 from the GEO databases. The intersection DEGs were analyzed for functional annotation and pathway enrichment. The GO and KEGG results showed that the DEGs were closely related to NOA immunity. We further identified and validated the hub genes among the DEGs. We found most hub genes were highly expressed in macrophages while they were low expressed in other immune cells. It indicated the importance of macrophages in the testicular microenvironment. AR was highly expressed in macrophages of AZFa_Del. The research found AR could promote or inhibit the inflammatory characteristics of mouse macrophages, depending on the external environmental condition, AR expression, and hormone concentration(14). About 75% of macrophages highly expressed DCN dan APOE, decorin (DCN) bound to the receptor on macrophages to trigger the production of pro-inflammatory cytokines (15). Interestingly, in iNOA patients, the expression of hub genes in macrophages was similar to normal adults, while in KS patients even down-regulated. But those hub genes were expressed in other immune cells obviously. This further indicated the importance of studying immune intercellular communication.

Next, we compared the immune cells in the microenvironment in normal subjects with that in NOA patients. We found that NOA patients had the largest difference in macrophages compared with normal adult patients. The number of macrophages is larger than that of normal adults and significantly reduced B cells. Under specific physiological conditions, macrophages pass through the epithelium of the tubules seminiferous as well as of the rete testis to phagocytose degenerating spermatozoa(16). Recent research suggested that the accumulation of infiltrating macrophages could impair normal testicular homeostasis by compromising the beneficial immunosuppressive environment of the organ(6). In our study, we found an abnormal increase of macrophages in patients. Because the development of Sertoli cells was impaired and the blood-testis-barrier weakened. Macrophages may be involved in the pathogenesis of azoospermia. The immune environment depends on the cooperation of multiple immune cells. We further analyzed the relationship between immune cells in the testicular microenvironment to provide clues for the treatment of NOA.

We used CellChat to analyze the relationship between the immune cells of normal subjects at the molecular level. Firstly, we found the great significance of macrophages and B cells in the immune environment from the size of nodes and the number of connections (**Figs 8A, and S1 Fig**). The MIF-(CD74+CXCR4) ligand-receptor pair has the highest contribution to immune cell interaction in infancy. NK cells receive signals from the other four kinds of immune cells through it. Macrophage migration inhibitory factor (MIF) is a multifunctional molecule that significantly decreased with age in animal experiments(17). CD74 has not only the function of antigen presentation but also the function(18), CXCR4 was higher in prepubertal calves when compared to adult bulls(19). All those indicated the importance of MIF-(CD74+CXCR4) in maintaining immune homeostasis in infancy. PTN signal pathway almost through the whole age. The contribution of PTN-NCL to the pathway can’t be replaced. Pleiotrophin (PTN) with multiple functions which are expressed by endothelial cells and macrophages(20, 21). Nucleolin is a multifunctional protein which highly expressed in lymphocytes, monocytes, and neutrophils in dogs with malignant neoplasia(22). purified NCL can activate monocytes and macrophages (23). But there are few studies on the effect of PTN-NCL on male spermatogenesis. A large number of experiments are needed to verify it later.

In our research, there were huge differences in immune cells in the testicular microenvironment between normal subjects and NOA patients. We also analyzed three kinds of azoospermia. Although there were a few B cells in the microenvironment in iNOA. It seemed that they have the same microenvironment as normal subjects, PTN pathway played an important role in its cell communication and PTN-NCL also brought the greatest contribution to the pathway. However, macrophages could only receive signals. In our analysis, we found that MDK-LRP1 and MDK-NCL, which contribute the most to the MK pathway, respectively played an important role in the communication of immune cells in the testicular microenvironment of AZFa_Del and KS. Macrophages in AZFa_Del received signals from themselves. While macrophages in KS played the role of the signal source and receive the signal from other immune cells. It has been proved that LRP1 is highly expressed in macrophages(24). The mice homozygote knocks out the LRP1 gene and died in early embryonic development, indicating that LRP1 played a key role in development(25). In tumor research, it was found that the increase of MDK led to the interaction of its receptor LRP1, which was expressed by tumor-infiltrating macrophages and promoted the differentiation of immunosuppressive macrophages(26). Combined with the significance of RNA-seq, scRNA-seq and immune cells of azoospermia showed different cellular signal communication from normal subjects. We hypothesized that PTN-NCL, MDK-LRP1, and MDK-NCL may be the key points of immune cells in iNOA, AZFa_Del, and KS respectively.

## Conclusion

In conclusion, our research showed that in the immune microenvironment of azoospermia testis, the distribution of immune cells has changed dramatically compared with that of normal subjects. The most typical was the massive infiltration of macrophages. Abnormal communications in NOA compared with normal subjects between macrophages and other immune cells. Finally, we analyzed and compared the interactions between immune cells in normal subjects and azoospermia patients. Found that PTN-NCL ligand-receptor played a key role in iNOA, MDK-LRP1 in AZFa_Del, and MDK-NCL in KS. Because of the destruction of the azoospermia blood-testis-barrier, macrophages may be overactivated through other immune cells and participate in the destruction of sperm. Our findings form the basis for future research. These biomarkers are of great significance and may be the targets of immunotherapy for azoospermia. Understanding their complex role in testis will help to understand the mechanism of azoospermia more comprehensively, and provide new ideas for immunotherapy of azoospermia.

## Materials and methods

### Data collection and processing

The GEO database (http://www.Ncbi.nlm.nih.gov/geo/) is a public database used to host high-throughput microarray and next-generation sequence functional genomic datasets(27). We downloaded expression profiles of patients with NOA from the GEO database. We selected the datasets GSE145467 and GSE9210. The data of GSE145467 were obtained with the GPL4133 Platforms (Agilent-014850 Whole Human Genome Microarray 4×44K G4112F) by Clinical Institute of Medical Genetics University Medical Centre Ljubljana and came from 10 showing obstructive azoospermia and 10 samples showing NOA. Similarly, the data of GSE9210 were based on the GPL887 Platforms (Agilent-012097 Human 1A Microarray (V2) G4110B) by Tokai University School of Medicine. We analyzed 47 NOA and 11 obstructive azoospermia. We also selected the dataset with accession number GSE149512. The samples were obtained from public databases and the study was carried out in accordance with relevant guidelines/regulations. The statement of ethics approval and informed consent were not needed for this study.

### Identification of DEGs

We used GEO2R to separately screen DEGs between NOA tissue samples and OA tissue samples from the GSE145467 and GSE9210 datasets. DEGs were defined using the threshold of fold change>1.5 and an adjusted P-value (adj.p.val) ≤ 0.05. the differential expression of the gene is judged to be statistically significant. Next, we used the R software to extract the common DEGs between the two datasets and visualized them using Volcano plots, and Venn plots(28)

### Gene ontology (GO) and Kyoto encyclopedia of genes and genomes (KEGG) analysis

GO(gene ontology) databases define biological processes (BP), cellular components (CC), and molecular functions (MF) based on gene products and are used widely to interpret genomes(29). The KEGG database links genomic and functional information, allowing users to analyze gene function(30). DAVID (https://David.ncifcrf.gov/) is a free online tool that extracts biological significance from large lists of genes or proteins, providing users with functional annotation and visualization(31). In this study, we used DAVID f to obtain GO functional enrichment analyses and enriched KEGG pathways of the DEGs. P<0.05 was considered statistically significant and the results were visualized using ggplot2 in R.

### Acquisition of hub genes by PPI network analysis

The STRING database (https://string-db.org/, version: 11.5) was used to predict PPI networks from DEGs and to analyze the interaction between proteins (32). We used the STRING website to construct the PPI network DEGs, and used a minimum interaction score of 0.4. The plug-in cytoHubba in Cytoscape (version3.9.0) was used to visualize the protein-protein interaction (PPI) network, and identified the central genes by maximal clique centrality (MCC, one of the 12 methods to explore important nodes in biological networks) computing method(33).

### Single-cell sequencing data processing

In this study, the original gene expression matrix from the dataset GSE149512 was used, included 10 normal patients, and 7 azoospermia patients which included 1 AZFa_DEL, 3 iNOA, and 3 KS. with the aim of applying scRNA-seq data to dissect the immune cell composition and molecular features of the testicular microenvironment in NOA. The quality control (QC) process used the R package Seurat (version 4.1.1). Data filtering indicators were as follows: 1. The number of genes is greater than 300. 2. Cells with RNA greater than 1000 and less than 20000. 3. Less than 12.5% mitochondrial genes. 4. Ribosomal genes accounted for more than 0.3%. 5. Less than 3% erythrocyte gene. At the end of QC, the cell populations were annotated according to marker genes, and markers were identified as major immune cell types and visualized with the dimensionality reduction algorithm t-distributed stochastic neighbor embedding (t-SNE)(34).

### Inference and analysis of intercellular communication using CellChat

ScRNA-seq data contains gene expression information that could be used to infer inter intercellular communication. CellChat is a tool that could quantitatively infer and analyze intercellular communication networks from scRNA-seq data(11). We applied CellChat analysis to the scRNA-seq data of the scRNA-seq data of the NOA and normal samples. CellChat contains a database of receptor-ligand interactions. To obtain more critical cell-cell interactions in the testicular microenvironment, we selected receptor-ligand pairs for further analysis, aiming to explore the potential interactions between immune cells

## Supporting information

**S1 Fig.**
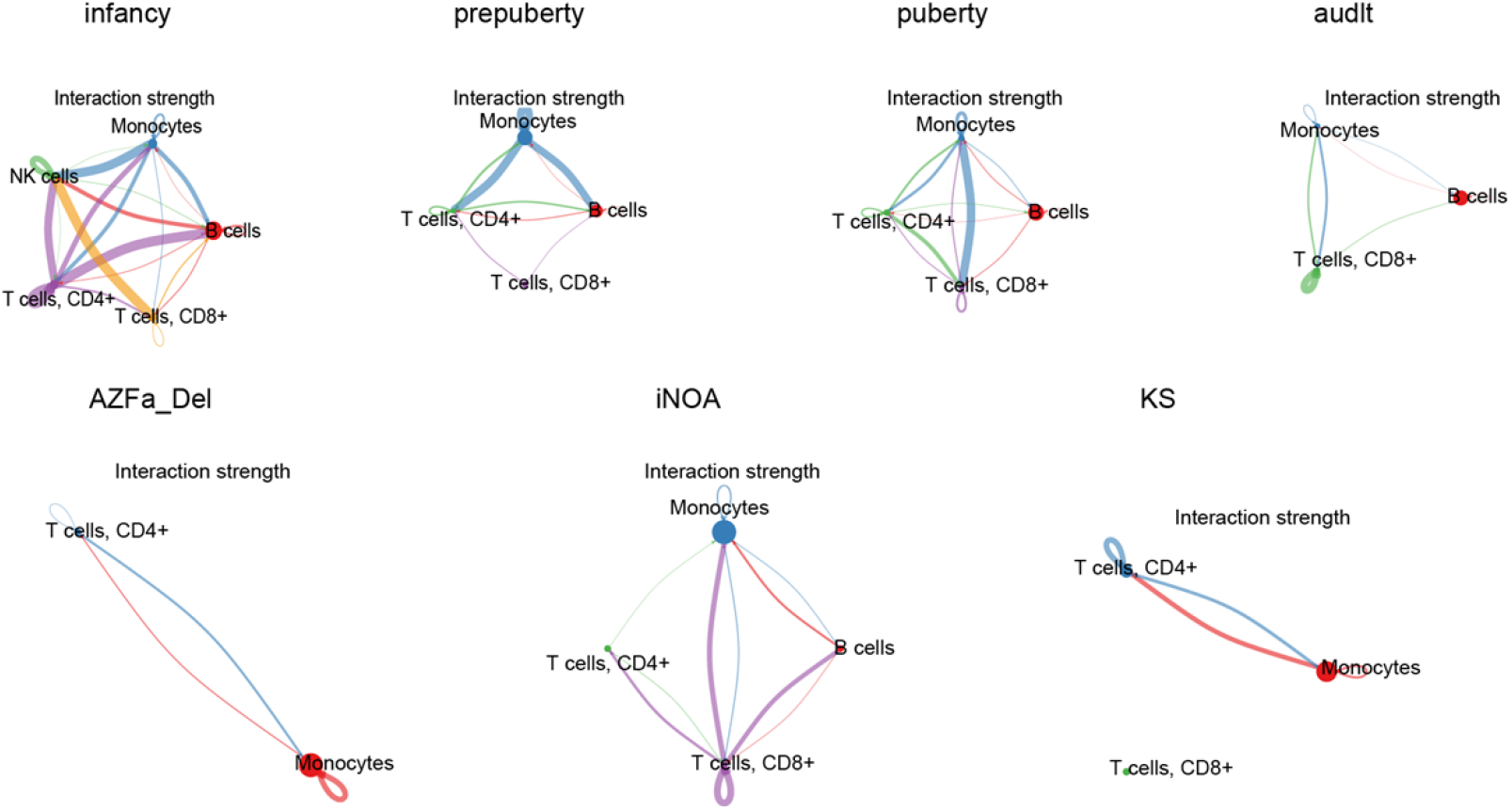
The strength of ligand-receptors interactions between different cell types. We compared the difference of the strength of ligand receptors between normal subjects and three kinds of azoospermia

## Author contributions

**Conceptualization:** YuanYuan Wu, Fang Wang

**Data curation:** YuanYuan Wu, JinGe Huang

**Formal analysis:** YuanYuan Wu, JinGe Huang, Nan Ding

**Funding acquisition:** Fang Wang

**Investigation:** MengHao Lu

**Methodology:** YuanYuan Wu, Fang Wang, MengHao Lu

**Project administration:** Fang Wang

**Resources:** Fang Wang

**Software:** YuanYuan Wu

**Supervision:** Fang Wang, Nan Ding

**Visualization:** YuanYuan Wu

**Writing-original draft:** YuanYuan Wu, Fang Wang, JinGe Huang, Nan Ding, MengHao Lu

**Writing-Review:** YuanYuan Wu, Fang Wang, JinGe Huang, Nan Ding, MengHao Lu

## Acknowledgments

We sincerely thank all participants in the study.

## Publisher’s note

All claims expressed in this article are solely those of the authors and do not necessarily represent those of their affiliated organizations, or those of the publisher, the editors, and the reviewers. Any product that may be evaluated in this article, or claim that may be made by its manufacturer, is not guaranteed or endorsed by the publisher.

## Data availability statement

Publicly available datasets were analyzed in this study. This data can be found here: http://www.Ncbi.nlm.nih.gov/geo/.

